# Identifying genetic variants associated with cerebellar volume in 33,265 individuals from the UK-Biobank

**DOI:** 10.1101/2020.11.24.393249

**Authors:** Tom Chambers, Valentina Escott-Price, Sophie Legge, Emily Baker, Krish D. Singh, James TR Walters, Xavier Caseras, Richard JL Anney

## Abstract

There is expanding interest in researching the cerebellum given accumulating evidence of its important contributions to cognitive and emotional functions, in addition to more established sensorimotor roles. While large genome-wide association studies (GWAS) have shed light on the common allele architecture of cortical and subcortical brain structures, the cerebellum remains under investigated. We conducted a meta-GWAS of cerebellar volume in 33,265 UK-Biobank European participants. Results show cerebellar volume to be moderately heritable (h^2^_SNP_=50.6%). We identified 33 independent genome-wide associated SNPs with total cerebellar volume, with 6 of these SNPs mapped to protein-coding genes and 5 more shown to alter cerebellar gene expression. We highlight 21 unique candidate genes for follow-up analysis. Cerebellar volume showed significant genetic correlation with brainstem, pallidum and thalamus volumes, but no significant correlations with neuropsychiatric phenotypes. Our results provide important new knowledge of the genetic architecture of cerebellar volume and its relationship with other brain phenotypes.

## Introduction

The cerebellum has historically been ascribed solely to a role in the coordination of movement, however, increasing evidence has underlined its relevance in cognition and emotional processing^1^. Detailed functional mapping of the cerebellum indicates expansive functional connectivity with non-motor cortical regions^2–4^ as well as elevated activity during a wide range of cognitive tasks^5^. Supporting its role in cognition, lesions and disruption of cerebellar functioning lead not only to motor alterations, but also to uncoordinated thought (i.e. dysmetria of thought)^6^, mirroring impairments present in some neurological and psychiatric disorders^7,8^.

Twin studies have estimated cerebellar volume to have moderate to high heritability (33.6 to 86.4%)^9^ in line with other structural brain phenotypes. Recent genome-wide association studies (GWAS) for cerebral anatomical phenotypes have revealed their highly polygenic nature, with a substantial contribution to heritability from common alleles (e.g. thalamus single nucleotide polymorphism (SNP)-based heritability h^2^_SNP_= 47%, cortical surface area h^2^_SNP_= 34%)^10–12^ and their shared genetic liabilities with brain-related phenotypes such as cognition or psychiatric disorders. Whilst two previous brain-wide GWAS studies have included cerebellar volumetric measures amongst other phenotypes investigated^13,14^, there has been little exploration and discussion of these cerebellar findings in terms of their relationship with other brain-based measures and functional consequences of genetic variants.

We report here a GWAS of total cerebellar grey matter volume in 33,265 participants from the UK-Biobank cohort^15^ - increasing the sample size by more than 10,000 participants from the largest GWAS to date including cerebellar measures^14^. We completed two independent GWAS analyses with approximately half the total sample in each, corresponding to two Magnetic Resonance Imaging (MRI) data releases from the UK-Biobank. We examined the replicability of our results between both GWASs, followed by a meta-analysis of both sets of results. We report on the genome-wide significant regions identified, including functional annotation and gene expression analysis to identify likely related genes, in addition to assessing the genetic overlap with other brain-based (e.g. cortical thickness) and brain-related (e.g. general cognitive ability) phenotypes. Our primary focus was on the genetic architecture of total cerebellar volume, however, we provided additional lobe-specific analyses based on primary, horizontal and posterolateral fissure separations. Our study expands our understanding of the influence of common genetic variants on brain anatomy and the shared genetic liability across different anatomical brain and cognitive/clinical phenotypes.

## Results

### GWAS analyses for phase 1 and phase 2 data releases

We processed and analysed two independent samples from the UK-Biobank corresponding to two consecutive data releases of brain imaging data, henceforth referred to as phase 1 and phase 2 (see Methods). A total of 17,818 participants from phase 1 (age mean[min,max]= 63[45,80]yrs, 53% female) and a total of 15,447 participants from phase 2 (age mean[min,max]= 65[48,81]yrs, 53% female) had genotype data which passed quality control and were included in their respective GWASs of cerebellar volume (supplementary table 1). Genotype quality control was performed on each phase separately. A total of 6,193,476 SNPs passed quality control and were common to both phases. Using conditional and joint analysis (COJO)^16^ on each phase of the GWAS results, we identified 6 independent genome-wide significant associated regions (containing index SNP p< 5×10^−8^; LD range SNPs r^2^ > 0.2 to index SNP) in the phase 1 GWAS, and 6 independent genome-wide significant associations in the phase 2 GWAS (Figure 1; supplementary table 2A & 2B).

### Between-phase results’ reliability and validity

We examined the direct replication, genetic correlation and the out-of-sample predictive ability of the identified variants for each phase. All but 1 of the 6 index SNPs observed in phase 1 were replicated in phase 2 (p< 0.0083 {0.05/6}), while all 6 of the index SNPs identified in phase 2 were replicated in phase 1 (p< 0.0083 {0.05/6}). Four index SNPs were genome-wide significant across both phases (Supplementary tables 2A & 2B).

**Figure 1:**
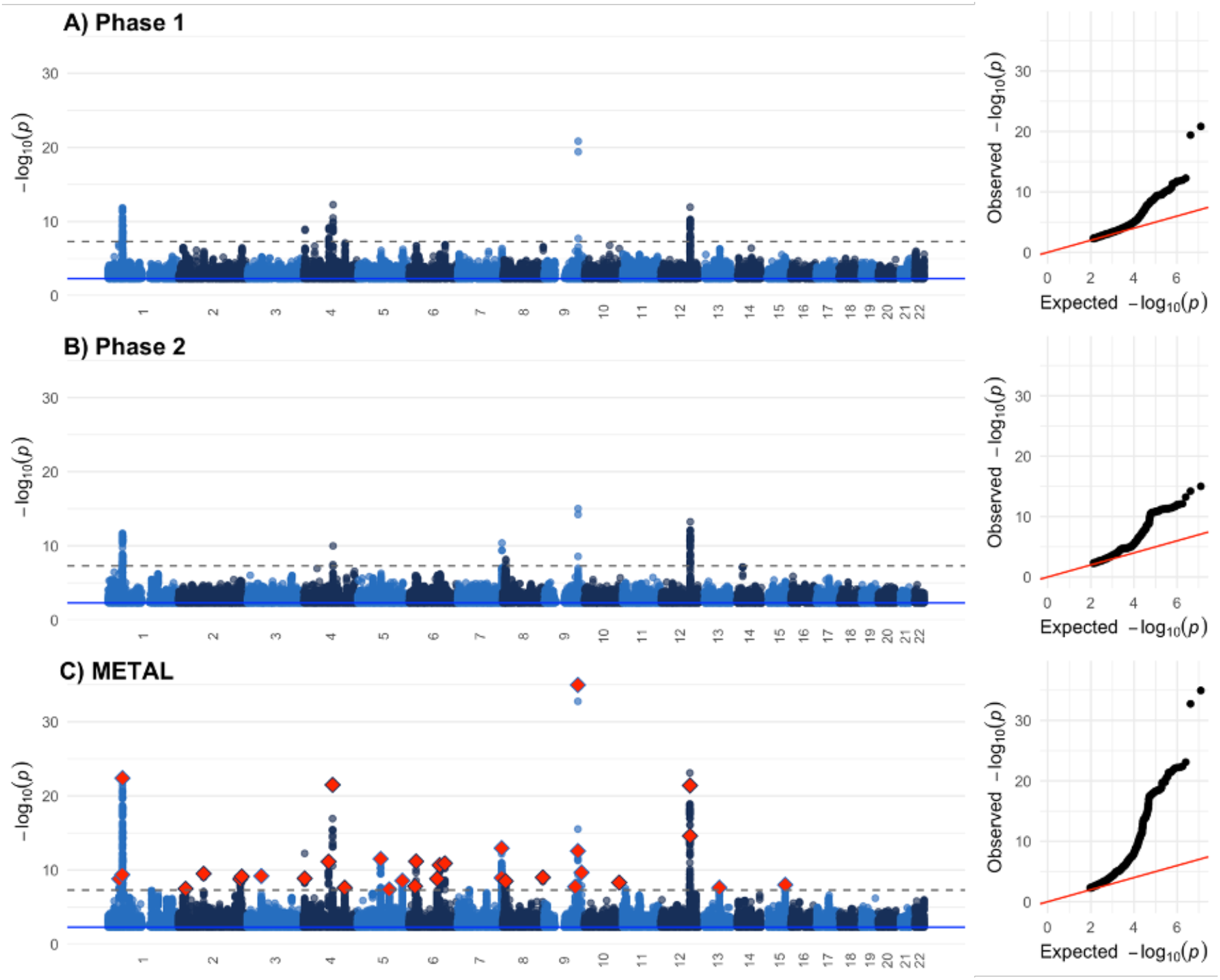
Manhattan plots of associations with total cerebellar volume for A) Phase 1 data release (n= 17,818), B) Phase 2 data release (n= 15,447), and C) Phase 1 + Phase 2 combined METAL meta-analysis. For the METAL plot, the 33 COJO identified independent index SNPs are highlighted (red diamond). In all cases, the dashed line indicates genome-wide significance at p < 5×10^−8^. Quantile-quantile (QQ) plots for each GWAS are provided next to the Manhattan plot. For all plots, points p > 5×10^−3^ (blue solid line) are removed for ease of interpretation.

We obtained similar moderate GCTA-GREML (Genome-wide complex trait analysis – Genome-based restricted maximum likelihood)^17,18^ SNP-based heritability for each phase (phase 1 h^2^_SNP_[standard error(SE)]= 46.8[3.4]% and phase 2 h^2^_SNP_[SE]= 45.3[3.9]%), and a very strong between-phase genetic correlation (*r*_*g*_[SE]= 1.0[0.1], p= 2.2×10^−33^).

Finally, the polygenic score based on the phase 1 GWAS that best predicted the variance of total cerebellar volume in phase 2 participants was that calculated at a SNP inclusion p-threshold< 0.01 (19,210 SNPs), uniquely explaining (ΔR^2^) 1.9% (p= 5.3×10^−118^) of the variance once the effect of the relevant demographic, imaging and genetic covariates (see methods) had been accounted for. Reciprocally, the polygenic score based on the phase 2 GWAS calculated at SNP inclusion p-0.1 (146,489 SNPs) uniquely accounted for the greatest amount of the variance of total cerebellar volume in phase 1 participants, being 1.3% (p= 3.9×10^−100^) (Supplementary table 3).

### Meta-analysis of GWAS results for phase 1 and phase 2

Given the high correlation between phases, we combined the summary statistics in a fixed-effect inverse-variance method meta-analysis using METAL^19^. The combined analysis included 33,265 participants and 6,193,476 SNPs present in both phases. The SNP-based heritability estimate in the combined sample was h^2^_SNP_[SE]= 50.6[2.0]%. Conditional analysis using COJO on the meta-GWAS summary statistics identified a total of 33 independent genome-wide significant associations (Figure 1; Table 1). LocusZoom^20^ figures of each of the 33 index SNPs are available in supplementary materials.

All index SNPs identified in the GWAS of each phase were included within the 33 independent SNPs identified in this meta-analysis; and all 33 index SNPs from this meta-analysis were nominally significant in each phase GWAS, with 32 and 29 of them remaining significant after Bonferroni correction (p< 0.0015 {0.05/33}) in the phase 1 and phase 2 GWAS, respectively.

**Table 1:**
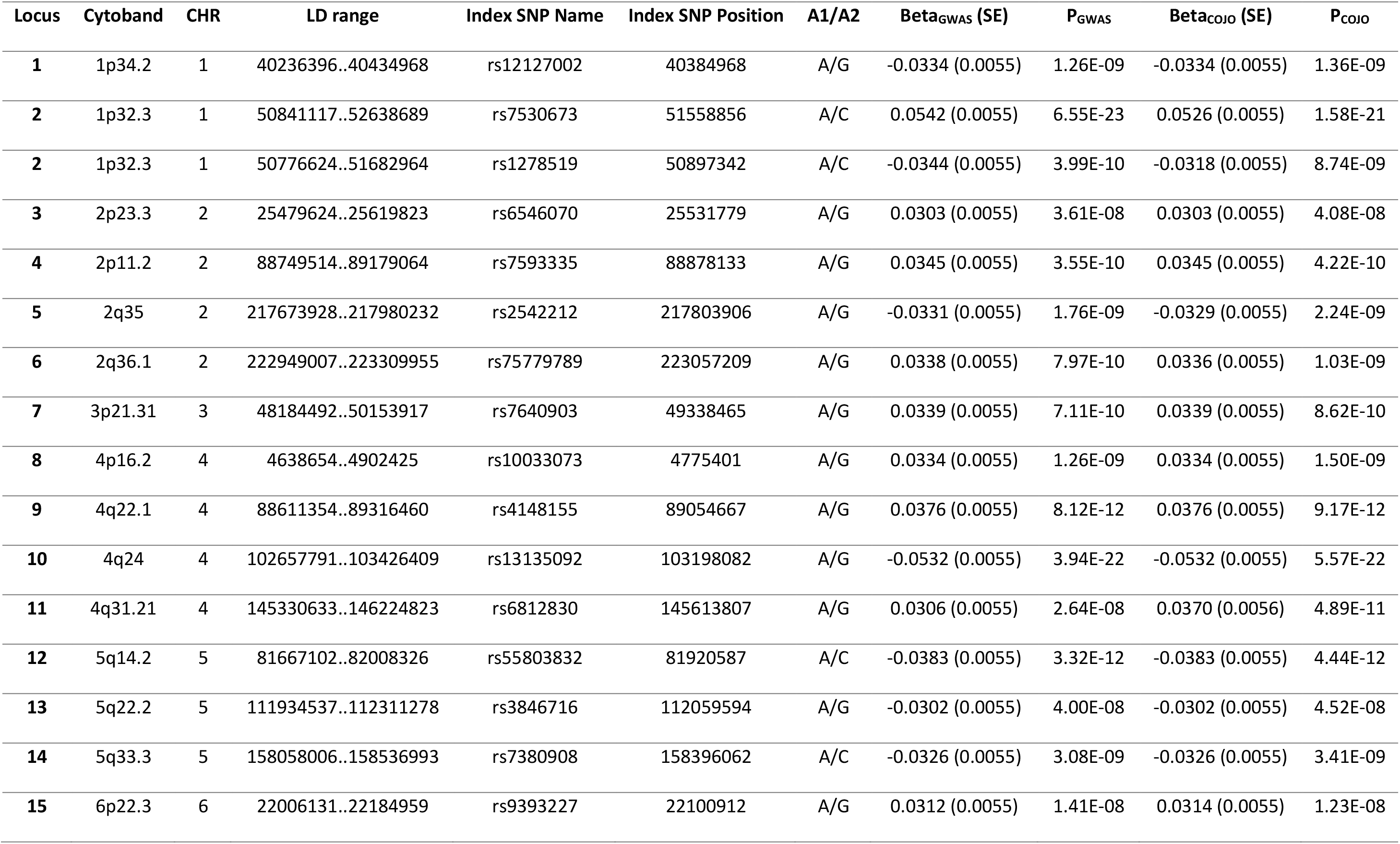
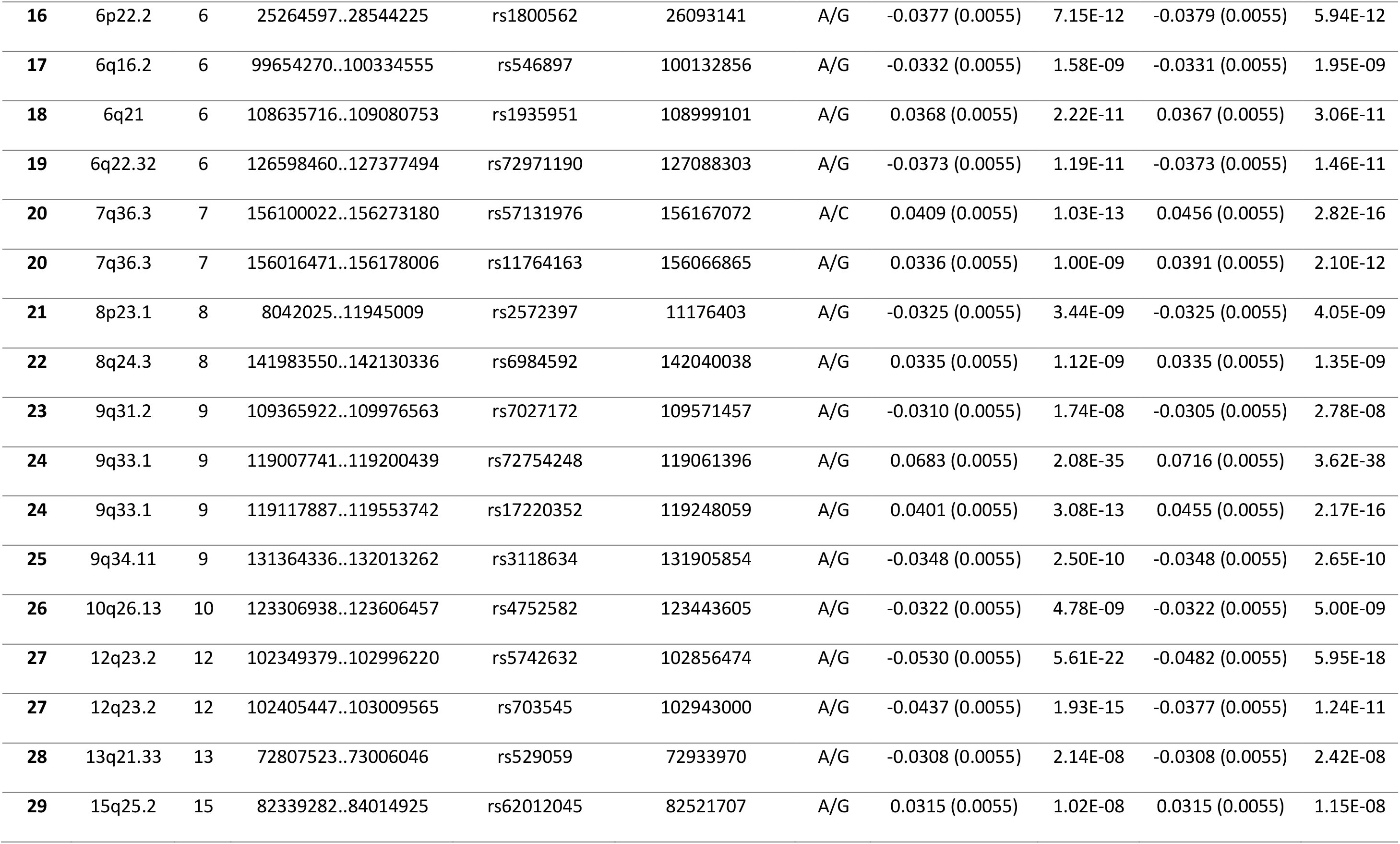
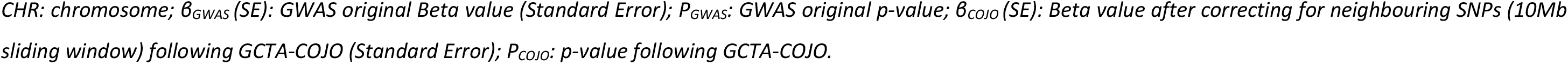
Genome-wide association results for total cerebellar volumes in European UK Biobank following COJO analysis.

| Locus | Cytoband | CHR | LD range | Index SNP Name | Index SNP Position | A1/A2 | Beta<sub>GWAS</sub> (SE) | P<sub>GWAS</sub> | Beta<sub>COJO</sub> (SE) | P<sub>COJO</sub> |
| --- | --- | --- | --- | --- | --- | --- | --- | --- | --- | --- |
| 1 | 1p34.2 | 1 | 40236396..40434968 | rs12127002 | 40384968 | A/G | -0.0334 (0.0055) | 1.26E-09 | -0.0334 (0.0055) | 1.36E-09 |
| 2 | 1p32.3 | 1 | 50841117..52638689 | rs7530673 | 51558856 | A/C | 0.0542 (0.0055) | 6.55E-23 | 0.0526 (0.0055) | 1.58E-21 |
| 2 | 1p32.3 | 1 | 50776624..51682964 | rs1278519 | 50897342 | A/C | -0.0344 (0.0055) | 3.99E-10 | -0.0318 (0.0055) | 8.74E-09 |
| 3 | 2p23.3 | 2 | 25479624..25619823 | rs6546070 | 25531779 | A/G | 0.0303 (0.0055) | 3.61E-08 | 0.0303 (0.0055) | 4.08E-08 |
| 4 | 2p11.2 | 2 | 88749514..89179064 | rs7593335 | 88878133 | A/G | 0.0345 (0.0055) | 3.55E-10 | 0.0345 (0.0055) | 4.22E-10 |
| 5 | 2q35 | 2 | 217673928..217980232 | rs2542212 | 217803906 | A/G | -0.0331 (0.0055) | 1.76E-09 | -0.0329 (0.0055) | 2.24E-09 |
| 6 | 2q36.1 | 2 | 222949007..223309955 | rs75779789 | 223057209 | A/G | 0.0338 (0.0055) | 7.97E-10 | 0.0336 (0.0055) | 1.03E-09 |
| 7 | 3p21.31 | 3 | 48184492..50153917 | rs7640903 | 49338465 | A/G | 0.0339 (0.0055) | 7.11E-10 | 0.0339 (0.0055) | 8.62E-10 |
| 8 | 4p16.2 | 4 | 4638654..4902425 | rs10033073 | 4775401 | A/G | 0.0334 (0.0055) | 1.26E-09 | 0.0334 (0.0055) | 1.50E-09 |
| 9 | 4q22.1 | 4 | 88611354..89316460 | rs4148155 | 89054667 | A/G | 0.0376 (0.0055) | 8.12E-12 | 0.0376 (0.0055) | 9.17E-12 |
| 10 | 4q24 | 4 | 102657791..103426409 | rs13135092 | 103198082 | A/G | -0.0532 (0.0055) | 3.94E-22 | -0.0532 (0.0055) | 5.57E-22 |
| 11 | 4q31.21 | 4 | 145330633..146224823 | rs6812830 | 145613807 | A/G | 0.0306 (0.0055) | 2.64E-08 | 0.0370 (0.0056) | 4.89E-11 |
| 12 | 5q14.2 | 5 | 81667102..82008326 | rs55803832 | 81920587 | A/C | -0.0383 (0.0055) | 3.32E-12 | -0.0383 (0.0055) | 4.44E-12 |
| 13 | 5q22.2 | 5 | 111934537..112311278 | rs3846716 | 112059594 | A/G | -0.0302 (0.0055) | 4.00E-08 | -0.0302 (0.0055) | 4.52E-08 |
| 14 | 5q33.3 | 5 | 158058006..158536993 | rs7380908 | 158396062 | A/C | -0.0326 (0.0055) | 3.08E-09 | -0.0326 (0.0055) | 3.41E-09 |
| 15 | 6p22.3 | 6 | 22006131..22184959 | rs9393227 | 22100912 | A/G | 0.0312 (0.0055) | 1.41E-08 | 0.0314 (0.0055) | 1.23E-08 |
| **16** | 6p22.2 | 6 | 25264597..28544225 | rs1800562 | 26093141 | A/G | -0.0377 (0.0055) | 7.15E-12 | -0.0379 (0.0055) | 5.94E-12 |
| **17** | 6q16.2 | 6 | 99654270..100334555 | rs546897 | 100132856 | A/G | -0.0332 (0.0055) | 1.58E-09 | -0.0331 (0.0055) | 1.95E-09 |
| **18** | 6q21 | 6 | 108635716..109080753 | rs1935951 | 108999101 | A/G | 0.0368 (0.0055) | 2.22E-11 | 0.0367 (0.0055) | 3.06E-11 |
| **19** | 6q22.32 | 6 | 126598460..127377494 | rs72971190 | 127088303 | A/G | -0.0373 (0.0055) | 1.19E-11 | -0.0373 (0.0055) | 1.46E-11 |
| **20** | 7q36.3 | 7 | 156100022..156273180 | rs57131976 | 156167072 | A/C | 0.0409 (0.0055) | 1.03E-13 | 0.0456 (0.0055) | 2.82E-16 |
| **20** | 7q36.3 | 7 | 156016471..156178006 | rs11764163 | 156066865 | A/G | 0.0336 (0.0055) | 1.00E-09 | 0.0391 (0.0055) | 2.10E-12 |
| **21** | 8p23.1 | 8 | 8042025..11945009 | rs2572397 | 11176403 | A/G | -0.0325 (0.0055) | 3.44E-09 | -0.0325 (0.0055) | 4.05E-09 |
| **22** | 8q24.3 | 8 | 141983550..142130336 | rs6984592 | 142040038 | A/G | 0.0335 (0.0055) | 1.12E-09 | 0.0335 (0.0055) | 1.35E-09 |
| **23** | 9q31.2 | 9 | 109365922..109976563 | rs7027172 | 109571457 | A/G | -0.0310 (0.0055) | 1.74E-08 | -0.0305 (0.0055) | 2.78E-08 |
| **24** | 9q33.1 | 9 | 119007741..119200439 | rs72754248 | 119061396 | A/G | 0.0683 (0.0055) | 2.08E-35 | 0.0716 (0.0055) | 3.62E-38 |
| **24** | 9q33.1 | 9 | 119117887..119553742 | rs17220352 | 119248059 | A/G | 0.0401 (0.0055) | 3.08E-13 | 0.0455 (0.0055) | 2.17E-16 |
| **25** | 9q34.11 | 9 | 131364336..132013262 | rs3118634 | 131905854 | A/G | -0.0348 (0.0055) | 2.50E-10 | -0.0348 (0.0055) | 2.65E-10 |
| **26** | 10q26.13 | 10 | 123306938..123606457 | rs4752582 | 123443605 | A/G | -0.0322 (0.0055) | 4.78E-09 | -0.0322 (0.0055) | 5.00E-09 |
| **27** | 12q23.2 | 12 | 102349379..102996220 | rs5742632 | 102856474 | A/G | -0.0530 (0.0055) | 5.61E-22 | -0.0482 (0.0055) | 5.95E-18 |
| **27** | 12q23.2 | 12 | 102405447..103009565 | rs703545 | 102943000 | A/G | -0.0437 (0.0055) | 1.93E-15 | -0.0377 (0.0055) | 1.24E-11 |
| **28** | 13q21.33 | 13 | 72807523..73006046 | rs529059 | 72933970 | A/G | -0.0308 (0.0055) | 2.14E-08 | -0.0308 (0.0055) | 2.42E-08 |
| **29** | 15q25.2 | 15 | 82339282..84014925 | rs62012045 | 82521707 | A/G | 0.0315 (0.0055) | 1.02E-08 | 0.0315 (0.0055) | 1.15E-08 |

*CHR: chromosome; $\beta_{\text{GWAS}}$ (SE): GWAS original Beta value (Standard Error); $P_{\text{GWAS}}$ : GWAS original p-value; $\beta_{\text{COJO}}$ (SE): Beta value after correcting for neighbouring SNPs (10Mb sliding window) following GCTA-COJO (Standard Error); $P_{\text{COJO}}$ : p-value following GCTA-COJO.*

### Annotation of genome-wide significant regions from the meta-GWAS

We identified high linkage disequilibrium (LD) partners (r^2^> 0.8) of the 33 index SNPs, mapped nearby genes and annotated all SNPs within each region with SNP Consequence, CADD Phred score, and putative functional consequence via PolyPhen and SIFT category (Supplementary tables 4, 5A & 5B). Five index SNPs were directly or in high LD (r^2^> 0.8) with non-synonymous SNPs, causing alterations in protein structure. Of these 5 non-synonymous SNPs, two were flagged as likely deleterious: the missense variant rs1800562 in the *HFE* (Hereditary hemochromatosis type 1) homeostatic iron regulator gene and rs13107325 located within the metal cation symporter SLC39A8 (Solute carrier family 39 member 8). The other three non-synonymous SNPs were flagged as tolerated/benign and reside within the transcription factor *EIF2AK3* (Eukaryotic translation initiation factor 2 alpha kinase 3), the protein phosphatase *PPP2R4 (*Protein Phosphatase 2 Phosphatase Activator; alias *PTPA)*, and the transcription factor *MYCL* (MYCL proto-oncogene). A synonymous annotated SNP was located within the *PAPPA* (Pregnancy-associated plasma protein A) gene, being also the region with the most significant association with cerebellar volume from our results.

### Expression quantitative trait loci (eQTL) and summary-data-based Mendelian randomisation (SMR) analysis

SNPs within each independent region were also mapped to cis-eQTL SNPs from the Genotype-Tissue Expression – version 7 (GTEx-7) cerebellum and cerebellar hemisphere labelled datasets. Six of the independent regions contained genome-wide significant eQTLs for either cerebellar labelled tissue, at cytobands 3p21.31, 5q14.2, 6q16.2, 8p23.1, 8q24.3 and 9q34.11 (Supplementary table 6A & 6B). The index SNP rs3118634 at 9q34.11 is an eQTL for 3 transcripts; *PPP2R4*, as well as two transcripts of unknown function, *RP11-247A12.2* and *RP11-247A12.7*. A region within the 3p21.31 cytoband included the variant rs3774800, a SNP which is in strong LD with the index SNP rs7640903 of that region (r^2^= 0.83) and is an eQTL with 5 transcripts: *AMT* (Aminomethyltransferase)*, CCDC71* (Coiled-Coil Domain Containing 71), *NCKIPSD* (NCK Interacting Protein With SH3 Domain), *WDR6* (WD Repeat Domain 6) and *GPX1* (Glutathione Peroxidase 1), with the latter only observed as an eQTL in the cerebellum labelled dataset while the other four were all observed as eQTLs in both cerebellar labelled datasets. The variant rs55803832 located at 5q14.2 is an eQTL in cerebellum labelled tissue for the extracellular matrix protein gene *VCAN* (Versican). Additional eQTLs were mapped for *PTK2* (Protein Tyrosine Kinase 2) and other transcripts of unknown function, namely *RP1-199J3.5, RP11-481A20.10*, *RP11-481A20.11* and *AF131216.5*.

We further extended the eQTL investigation by applying SMR^21,22^ analysis (Table 2). SMR examines the relationship between the GWAS and eQTL association at multiple SNPs within a region and, by doing so, can distinguish between associations driven by linkage from those by possible causal (or pleiotropic) relationships between altered gene expression and altered cerebellar volume. We again focused our analysis on the two cerebellar labelled GTEx-v7 eQTL tissue datasets. SMR identified significant relationships between associations at 3 independent regions: at 5q14.2, 8p23.1 and 9q34.11 cytobands. In total there were 6 transcripts which showed evidence supporting a causal (or pleiotropic) relationship between trait association and transcript expression, namely *PPP2R4, RP11-247A12.2* and *RP11-247A12.7* at 9q34.11; *VCAN* at 5q14.2; and the long non-coding RNA *FAM85B* (family with sequence similarity 85 member B) and pseudogene *FAM86B3P* (family with sequence similarity 86 member B3, pseudogene) at 8p23.1. The strongest SMR association was observed for *VCAN*, where we see a clear relationship with the GWAS associations with cerebellar volume and *VCAN* gene expression in cerebellum labelled tissue (Figure 2). It is important to note that the physical location of the *VCAN* gene transcript does not overlap with the location of the peaks of the GWAS associations and that this relationship would not have been prioritised without the use of functional annotations.

**Table 2:** The number of genes identified by summary data-based Mendelian randomisation (SMR) analysis.

| Locus | Cytoband | Tissue | Probe ID | Gene Symbol | Top SMR Marker | Top SMR Marker Position | P (eQTL) | P (GWAS) | P (SMR) | P (HEIDI) | N SNPs HEIDI |
| --- | --- | --- | --- | --- | --- | --- | --- | --- | --- | --- | --- |
| 12 | 5q14.2 | Cerebellum | ENSG00000038427.11 | VCAN | rs55803832 | 81920587 | 1.48E-12 | 3.09E-12 | 6.93E-07 | 0.57 | 10 |
| 21 | 8p23.1 | Cerebellum | ENSG000000253893.2 | FAM85B | rs2980439 | 8094870 | 3.58E-21 | 1.01E-06 | 1.40E-05 | 0.43 | 20 |
| 21 | 8p23.1 | Cerebellar Hemisphere | ENSG000000173295.3 | FAM86B3P | rs1878561 | 8092405 | 2.85E-19 | 1.77E-06 | 2.44E-05 | 0.39 | 20 |
| 21 | 8p23.1 | Cerebellum | ENSG000000173295.3 | FAM86B3P | rs1878561 | 8092405 | 2.37E-25 | 1.77E-06 | 1.39E-05 | 0.12 | 20 |
| 25 | 9q34.11 | Cerebellum | ENSG000000119383.15 | PPP2R4 | rs3118634 | 131905854 | 3.99E-16 | 2.14E-10 | 5.87E-07 | 0.27 | 14 |
| 25 | 9q34.11 | Cerebellum | ENSG000000204055.4 | RP11-247A12.2 | rs3118634 | 131905854 | 6.18E-09 | 2.14E-10 | 1.87E-05 | 0.47 | 13 |
| 25 | 9q34.11 | Cerebellar Hemisphere | ENSG000000268707.1 | RP11-247A12.7 | rs3124505 | 131887856 | 1.94E-20 | 1.31E-08 | 1.31E-06 | 0.17 | 19 |
| 25 | 9q34.11 | Cerebellum | ENSG000000268707.1 | RP11-247A12.7 | rs3118634 | 131905854 | 1.16E-20 | 2.14E-10 | 1.65E-07 | 0.23 | 19 |

*P (eQTL/GWAS/SMR): p-values from the GWAS results, eQTL association, and SMR mediation tests; P (HEIDI): p-values from the HEIDI (heterogeneity in dependent instruments) test with $p > 0.05$ indicating pleiotropic (over linkage) associations; N SNPs HEIDI: number of SNPs used included in the HEIDI test*

**Figure 2:**
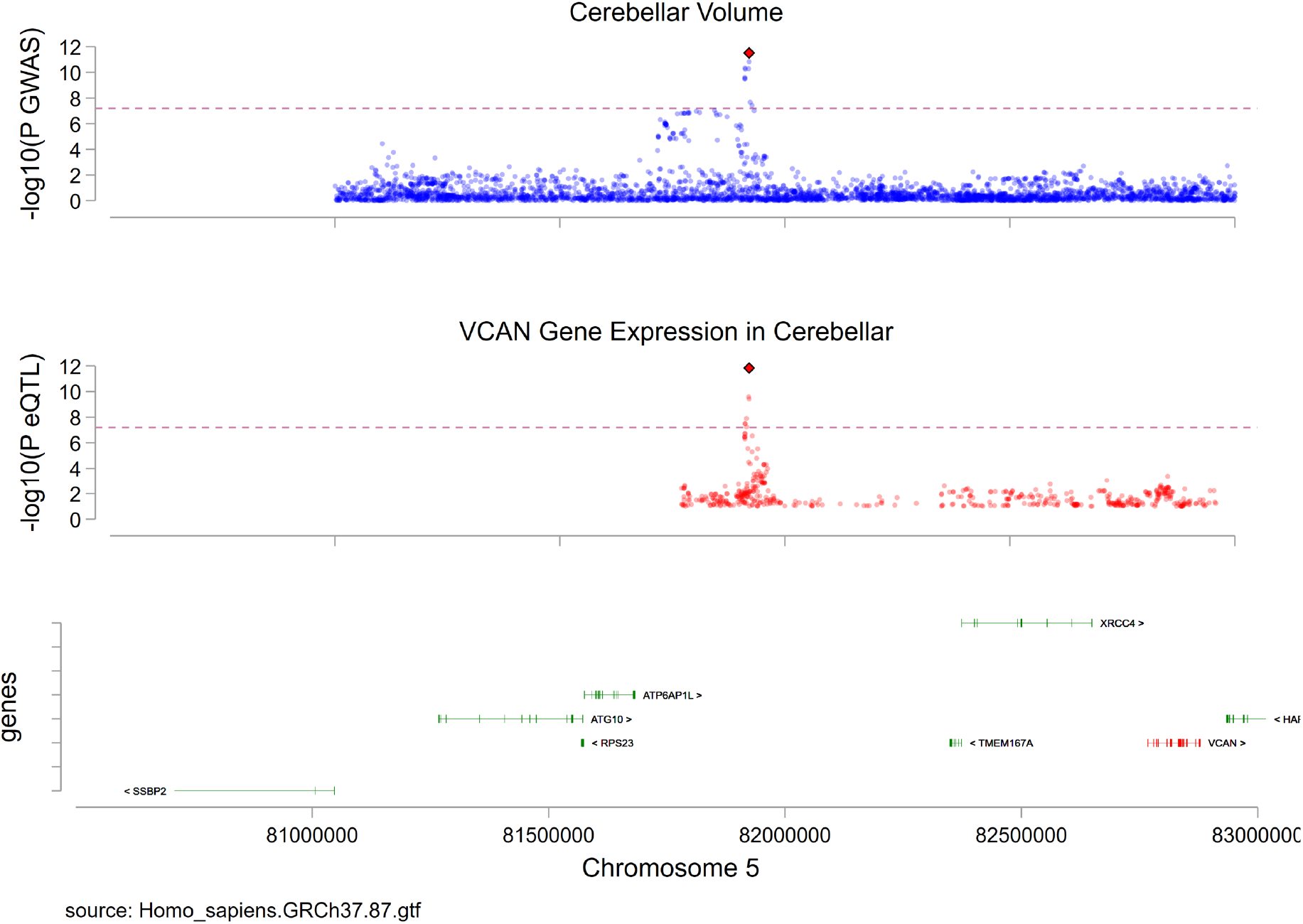
Prioritisation of *VCAN* gene transcript using SMR analysis. For the identified region, we show an association between SNP effect p-values from the meta-GWAS on total bran volume (top) and SNP effects on *VCAN* eQTL expression in GTEx-v7 cerebellum labelled tissue (middle) for all SNPs present in each summary data, in addition to gene transcripts present in that region (bottom). The highlighted red diamonds reflect the top SNP identified in each analysis which, in this instance, was the identified GWAS index SNP rs55803832.

### Genetic Correlations

#### Correlation against previous research on cerebellar phenotypes

We performed genetic correlation analyses between our meta-GWAS results and previous studies including cerebellar volumes: Elliott et al (2018)^13^ (left & right cerebellum) and Zhao et al (2019)^14^ (left & right cerebellar hemispheres and 3 vermal divisions). We found high genetic correlation between our results and those of Elliott et al (left and right cerebellum: r_g_[95% Confidence Intervals(CI)]= 0.92[0.75,1.00] & 0.98[0.77,1.00], respectively) and Zhao et al (left & right hemispheres; IIV-V, VI-VII & VIII-IX vermal regions: r_g_[95%CI]= 0.91[0.84,0.97] & 0.91[0.84,0.98]; 0.44[0.28,0.60], 0.45[0.32,0.57] & 0.56[0.46,0.65], respectively), with all passing Bonferroni corrected significance threshold (p< 0.0071 {0.05/7}) (Supplementary table 7A). Of the 33 independent associations that we identified, 15 were present in these previous works (our index SNP r^2^> 0.1 or the LD region around our index SNP< 500kb away from their identified independent regions) while 18 were novel to the literature.

#### Correlation against anthropomorphic phenotypes

Since several of the identified index variants were previously reported to be associated with multiple anthropometric traits (http://www.nealelab.is/uk-biobank/), as an additional analysis we investigated whether the variance in cerebellar volume observed might reflect measures of general body size. To do this, we explored genetic correlations between our cerebellar volume meta-GWAS and several anthropometric measurements in the UK-Biobank data, including Birth Weight, Body Fat Percentage, Body Mass Index (BMI), Sitting Height, Standing Height and Weight (Supplementary table 7B). None of these correlations were significant after Bonferroni correction (p< 0.0083 {0.05/6}). The strongest correlation observed was with Body Mass Index (r_g_[95%CI]= ȡ0.07[−0.12,−0.02], p= 0.01).

#### Correlation against other brain-based and brain-related phenotypes

We examined the genetic correlation between our meta-GWAS for cerebellar volume and the most recent GWAS for subcortical volumes^11,12^ and cortical thickness and surface area^10^. We found positive genetic correlations passing our Bonferroni corrected significance threshold (p< 0.005 {0.05/10}) between the volumes of the cerebellum and the volume of brainstem (r_g_[95%CI]= 0.47[0.37,0.58], p= 1.0×10^−18^), pallidum (r_g_[95%CI]= 0.31[0.19,0.43], p= 4.5×10^−7^) and thalamus (r_g_[95%CI]= 0.24[0.12,0.36], p= 6.5×10^−5^). A trend towards a negative correlation with cerebral cortical surface area was also found, but this just felt short of the Bonferroni corrected significant threshold (r_g_[95%CI]= −0.14[−0.25,−0.04], p= 0.007) (Table 3A).

We also ascertained the genetic correlation between cerebellar volume and brain-related phenotypes previously associated with cerebellar anatomy and/or function, including schizophrenia^23^, bipolar^24^, autism spectrum^25^ disorders, Parkinson’s disease^26^ and general cognitive ability^27^. None of these showed significant genetic correlation with cerebellar volume even at a nominal level of significance (p< 0.05) (Table 3B).

**Table 3:** Genetic correlation of total cerebellar volume with (A) brain-based phenotypes and (B) brain-related phenotypes previously associated with cerebellar anatomy/function.

| | $h^2_{\text{SNP}}$ (%) | $h^2_{\text{SNP}}$ SE (%) | $r_g$ | 95% Confidence intervals | | p | $p_{\text{Bonferroni}}$ |
| --- | --- | --- | --- | --- | --- | --- | --- |
| A) Brain-based phenotypes |  |  |  |  |  |  |  |
| Brainstem | 31.7 | 3.4 | 0.47 | 0.37 | 0.58 | 1.02E-18 | 1.02E-17 |
| Pallidum | 16.9 | 2.3 | 0.31 | 0.19 | 0.43 | 0.00000045 | 0.0000045 |
| Thalamus | 16.0 | 2.1 | 0.24 | 0.12 | 0.36 | 0.0000645 | 0.000645 |
| Cortical surface area | 35.3 | 3.2 | -0.14 | -0.25 | -0.04 | 0.007 | 0.07 |
| Amygdala | 8.4 | 1.9 | -0.18 | -0.37 | 0.01 | 0.07 | 0.67 |
| Hippocampus | 13.0 | 2.7 | -0.14 | -0.29 | 0.02 | 0.08 | 0.84 |
| Caudate | 28.6 | 2.6 | -0.07 | -0.18 | 0.04 | 0.20 | 1.00 |
| Accumbens | 20.2 | 2.3 | -0.07 | -0.20 | 0.06 | 0.29 | 1.00 |
| Putamen | 28.6 | 2.8 | 0.01 | -0.10 | 0.11 | 0.88 | 1.00 |
| Cortical thickness | 26.5 | 2.2 | -0.01 | -0.11 | 0.10 | 0.91 | 1.00 |
| B) Brain related phenotypes |  |  |  |  |  |  |  |
| Schizophrenia disorder | 42.1 | 1.5 | -0.04 | -0.10 | 0.02 | 0.18 | 1.00 |
| Bipolar disorder | 34.6 | 1.9 | -0.04 | -0.12 | 0.04 | 0.33 | 1.00 |
| Autism spectrum disorder | 19.5 | 1.5 | -0.10 | -0.22 | 0.02 | 0.10 | 0.62 |
| Cognition | 20.0 | 0.7 | 0.00 | -0.06 | 0.06 | 0.97 | 1.00 |
| Educational attainment | 11.2 | 0.3 | -0.02 | -0.07 | 0.03 | 0.43 | 1.00 |
| Parkinson's disease | 7.4 | 0.8 | 0.00 | -0.11 | 0.11 | 1.00 | 1.00 |

*Calculated using LDSC regression analysis software. $h^2_{SNP}$ : SNP-based heritability estimates; SE: standard error; $r_g$ : genetic correlation; $p$ : uncorrected $p$ -values; $p_{Bonferroni}$ : $p$ -values adjusted for the number of tests performed regions/traits tested (10 & 6, respectively)*

### Cerebellar lobe analysis

To ascertain the homogeneity of common allele architecture across the cerebellum, we partitioned the cerebellum into 7 separate lobes using the demarcations of primary, horizontal, and posterolateral fissures: hemispheres of the anterior (I-V), superior posterior (VI-Crus I), inferior posterior (Crus II-IX) and flocculonodular (X) cerebellum, plus the vermal regions of the latter three. We showed similar SNP-based heritability estimates across all lobes ranging around the overall cerebellar heritability, except for the vermal flocculonodular lobe which showed slightly lower heritability (h^2^_SNP_[SE]= 35.4[1.9]%) (Supplementary table 8A). Genetic correlation between lobes was at least moderate for most (between-lobes mean r_g_≈ 0.44) and all correlations survived Bonferroni correction for the total number of lobe-pairings tested (p< 0.0024 {0.05/21}); being strongest between the inferior posterior hemisphere and vermis (r_g_[95%CI]= 0.66[0.60,0.72], p= 1.4×10^−103^) and weakest between the flocculonodular hemisphere and vermis (r_g_[95%CI]= 0.19[0.07,0.30], p= 1.3×10^−3^) (Figure 3; Supplementary table 8A).

**Figure 3:**
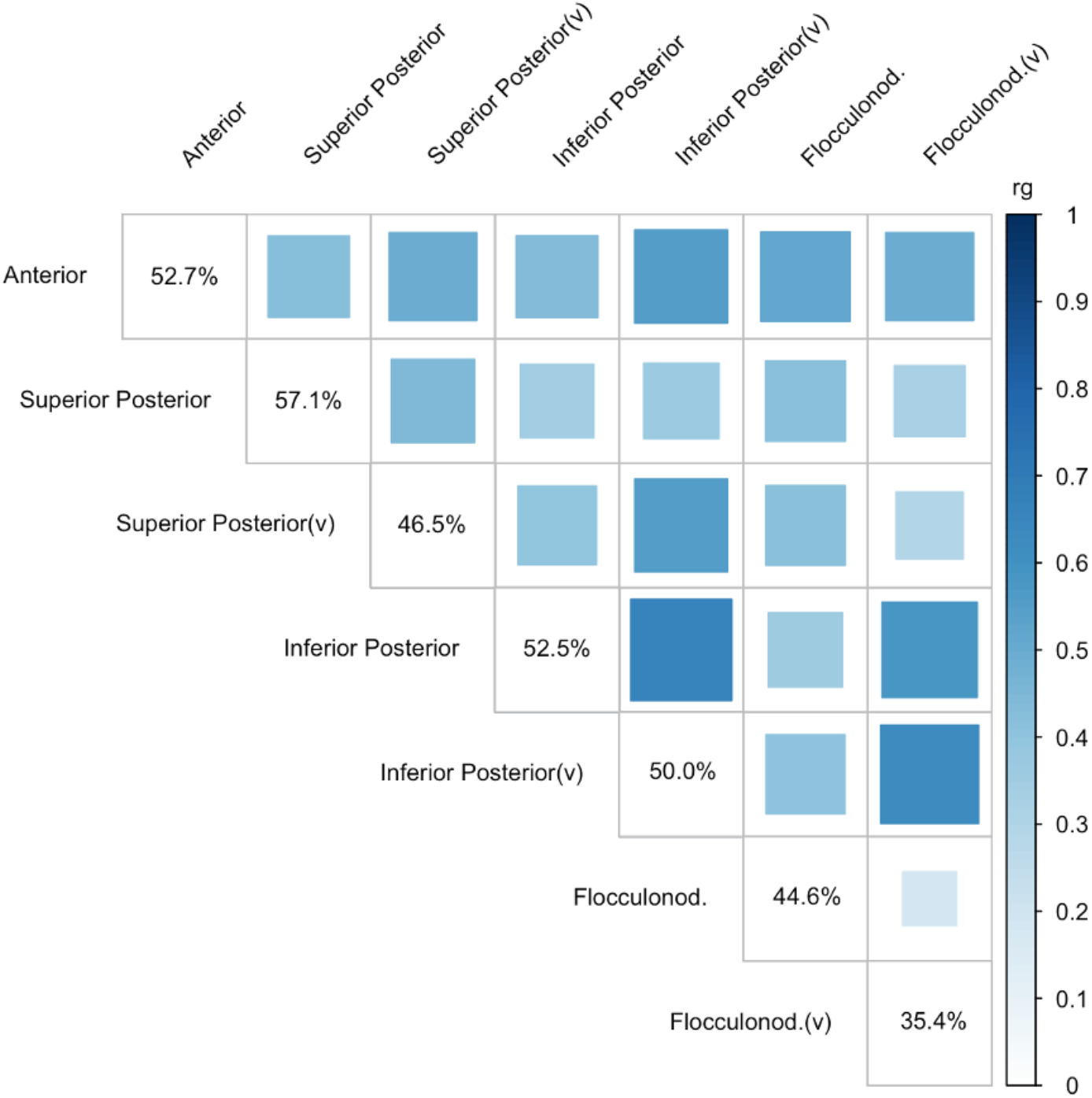
Genetic correlation between the seven cerebellar lobes. Tile size and colour represent genetic correlation values (r_g_) between lobes calculated using LDSC regression analysis. Diagonal values of SNP-based heritability estimates calculated using GCTA-GREML. All correlations passed Bonferroni correction p < 0.0024 {0.05/21}. (v): vermis.

We explored the associations between these cerebellar lobes and the brain-based and brain-related phenotypes mentioned previously. Correcting for the 70 possible pairings with other brain-based traits (p< 0.00071 {0.05/70}), positive correlations were seen between all lobes and the brainstem, being highest with flocculonodular vermis (r_g_[95%CI]= 0.49[0.38,0.59], p= 4.3×10^−18^) and lowest with the superior posterior hemisphere and vermis (r_g_[95%CI]= 0.26[0.15,0.37], p= 3.0×10^−6^ & r_g_[95%CI]=0.26[0.14,0.38], p= 1.5×10^−5^, respectively). All lobes also showed a positive genetic correlation with bilateral pallidum, aside from the superior posterior hemisphere which did not survive Bonferroni correction (r_g_[95%CI]= 0.18[0.06,0.30], p= 3.8×10^−3^) and with the highest correlation being with the flocculonodular vermis (r_g_[95%CI]= 0.30[0.17,0.43], p= 4.0×10^−6^). The same pattern was seen with the thalamus, with only the superior posterior hemisphere and vermis positive correlations not surviving Bonferroni correction (r_g_[95%CI]= 0.12[0.00,0.24], p= 0.066 & r_g_[95%CI]= 0.15[0.03,0.27], p=0.016, respectively) and with the highest correlation being with the flocculonodular vermis (r_g_[95%CI]= 0.31[0.18,0.45], p= 3.8×10^−6^) (Supplementary table 8B). We also found a negative genetic correlation between the flocculonodular hemispheres and cerebral cortical surface area (r_g_[95%CI]= −0.16[−0.26,−0.07], p= 4.2×10^−4^). Finally, no lobes showed Bonferroni-corrected significant (p< 0.0012 {0.05/42}) genetic correlation with any of the brain-related phenotypes included in our study (Supplementary table 8C).

## Discussion

In this study we combine the UK-Biobank imaging and genotype data of 33,265 individuals of European ancestry to investigate common allele influences on cerebellar volume. After ascertaining that total cerebellar volume was moderately heritable in our sample (h^2^_SNP_= 50.6%), we identified 33 independent genome-wide significant SNPs across 29 regions associated with this phenotype. Functional annotation and positional mapping identified 6 SNPs impacting protein coding genes while, via SMR, we show evidence of impact on expression of 6 transcripts in cerebellar tissue. Overall, we identified 21 genes of interest for follow-up analysis for their effect on cerebellar volume. We found a large genetic overlap between cerebellar volume and the volume of the brainstem, the pallidum and the thalamus, however, no genetic associations with neurological, psychiatric, or cognitive phenotypes previously associated with changes in cerebellar anatomy and/or function were found. Further analyses separating the cerebellum into lobes showed moderate to high genetic correlation between them, consistent with the relatively homogenous gene expression seen across cerebellar subdivisions^28^.

We initially performed two independent GWASs of cerebellar volume (phase 1 and phase 2) following two consecutive brain imaging data releases from the UK-Biobank. We obtained a high replication of independent index SNPs across phases, a significant out-of-sample prediction of cerebellar volume for both sets of results and a very high significant correlation between both GWASs. On this basis, we combined both sets of results into a meta-analysis to increase statistical power to reveal significant associations and for additional downstream investigation. We compared the main results from our meta-analysis to those previously reported on cerebellar grey-matter measures. To our knowledge only two previous GWASs have considered the cerebellum; both using UK-Biobank samples including approximately 10,000^13^ and 20,000^14^ participants each. We found high genetic correlation between our results for overall cerebellar volume and those previously reported on the left and right cerebellar hemispheres (including splitting of vermal regions in one^13^) in both these studies (all r_g_> 0.90). We found only moderate, although significant, correlation with those reported for purely vermal regions^14^ (mean r_g_≈ 0.50), with the reduction in genetic correlation likely due to their smaller volumes and so contributing less to the overall total cerebellar volume measure. Furthermore, the SNP-based heritability estimates we obtained are in keeping with those previously reported for other non-cerebellar grey-matter volumes^10–14^. Finally, since several of the independent genome-wide significant SNPs we identify had also previously been shown to be associated with multiple anthropometric traits (http://www.nealelab.is/uk-biobank/) – in addition to other brain-based and brain-related traits – we sought to confirm that our results were not simply a function of these anthropomorphic measures. We found no genetic correlation between our GWAS results for cerebellar volume and previous GWASs of anthropomorphic measures including birth weight, body fat percentage, body mass index (BMI), sitting height, standing height and weight. All of the above provide confidence about the reliability and validity of the results reported here.

We applied Conditional and Joint Analysis of Association (COJO) to the total cerebellar volume GWAS and identified 33 independent genome-wide significant SNP associations across 29 loci. Of the 33 independent SNPs, 15 had been directly or indirectly identified as showing association with cerebellar volume while 18 were novel^13,14^. One previously implicated SNP was the synonymous SNP rs35565319 in the *PAPPA* gene transcript. *PAPPA* is an IGF binging protein protease with possible cerebellar-specific interactional effects^29^, being highly expressed in the placenta and whose reduced protein expression is associated with various adverse pregnancy outcomes^30,31^ and neuronal survival in animal models^32^. Of the novel independent regions, 5 contained non-synonymous SNPs altering protein structure. Based on functional annotations, two of these were deleterious missense variants: rs13107325 in the metal cation symporter *SLC39A8* and rs1800562 in the homeostatic iron regulator *HFE.* The rs13107325 variant has been previously associated with increased volume in individual inferior posterior and flocculonodular lobules^13^, as well as with increased striatum and putamen volumes^13^. A study of the association between rs13107325 and putamen volume found it to be accompanied by decreased *SLC39A8* expression in the putamen and with the SNP-trait association decreased in those with schizophrenia^33^. The rs13107325 SNP has also been associated with schizophrenia itself^23^, neurodevelopmental outcomes and intelligence test performance^34,35^, blood pressure^36^ and numerous other factors^13,37,38^, including over 70 anthropometric traits (http://www.nealelab.is/uk-biobank/). The rs1800562 *HFE* SNP is also known as Cys282Tyr and has been associated with reduced putamen volume and T2star signal in the striatum^13^, as well as being involved in iron regulation and transport, being a major risk variant for hemochromatosis where it accounts for approximately 85% of cases^39^, mineral metabolism and haematological disorders^40^ ^13^. The other three non-synonymous SNPs included variants altering protein structure of translation initiation factor kinase (rs867529 in *EIF2AK3),* protein phosphatase (rs2480452 in *PPP2R4*) and proto-oncogene transcription factor (rs3134614 in *MYCL*) proteins.

Using eQTL cerebellar tissue data, we were also able to link SNPs within 6 of our associated regions with altered expression of 14 gene transcripts: *AF131216.5, AMT, CCDC71, GPX1, NCKIPSD, PPP2R4, PTK2, RP1-199J3.5, RP11-247A12.2, RP11-247A12.7, RP11-481A20.10, RP11-481A20.11, VCAN,* and *WDR6.* Use of summary-data-based Mendelian Randomisation (SMR) highlighted possible mediation effects of gene expression on SNP-trait associations for six gene transcripts at 3 of our associated regions: *PPP2R4*, *RP11-247A12.2* and *RP11-247A12.7; VCAN*; and pseudogenes *FAM86B3P* & *FAM85B. PPP2R4/PTPA*, therefore, was identified in both the functional annotation, eQTL-only and SMR follow-up analyses. Located at 9q34.11, *PPP2R4* encodes an activator of phosphatase 2A implicated in controlling cell growth and division, it has been shown to be expressed in neurones and glia in the brain, including the cerebellum, where it plays a role in regulating dendritic spine morphology^41^ and whose dysfunction is a known cause of spinocerebellar ataxia^42^. The strongest SMR association was with *VCAN*, which encodes the extracellular matrix protein Versican and which plays a number of crucial roles in maintaining the extracellular matrix, including in nervous system development^43,44^. The pseudogenes *FAM86B3P* and *FAM85B* were identified from the SMR analysis. *FAM85B*, as well as the other non-coding gene eQTLs for *RP11-481A20.10 and RP11-481A20.11* in the same region, have been indicated in mood instability and schizophrenia^45,46^. While a higher confidence can be placed on genes identified in SMR analyses, its requirement for multiple eQTL signals means that it also might omit genes with poorer coverage, therefore, both eQTL-only and SMR identified genes should be considered for future follow-up work.

In total, 732 unique gene transcripts were located within 500kb of the 33 independent genome-wide associated SNPs. Using functional annotations and gene expression data in the cerebellum, we refined this to a list of 21 gene transcripts which particularly warrant further interrogation and follow-up analysis due to our tagging of their protein coding regions or altered expression in cerebellar tissue (Supplementary table 9).

We found strong genetic correlation between the results of our cerebellar volume GWAS and those previously run on the volume of the brainstem, pallidum and thalamus, but not with any other subcortical structure or with cortical surface area or thickness. A clustering of genetic correlations between pallidum, thalamus and brainstem had been noted previoulsy^11^, as well as basal ganglia-thalamic pairings in twin-based imaging studies^47^. These results indicate a significant sharing of common allele influences on the volume of these four brain structures. This is at odds, however, with the correlations of their actual volumes, where significant (phenotypic) correlations are found across all subcortical volumes and with no particular clustering of the pallidum, thalamus and brainstem^11^. The genetic clustering of the cerebellum with these three subcortical structures might be explained by their white matter connectivity within the brain, particularly since the gene expression profile of cerebellar grey matter is quite distinct^28^. The major input and output nuclei of the cerebellum are located within the brainstem and thalamus, respectively, and the interaction between the pallidum and the cerebellum is also well known, occurring at the level of cortex, at the ventrolateral thalamus and/or via direct connections^48–50^. Both structures share roles in sensorimotor regulation, adaptation, learning and reward^48^. The common allele overlap correlation found across these four brain structures warrants further research into the neurobiological underpinnings of this potential network.

Perhaps surprisingly, considering the phenotypic association between grey matter volume in the cerebellum with cognitive function and psychopathology^51–53^, we did not find any evidence of a significant genetic correlation between cerebellar volume and our list of cognitive/neuropsychiatric phenotypes. Notably, previous GWASs of other brain-based phenotypes have also generally reported a lack of genetic association with most of these brain-related traits despite clinical research showing brain-wide anatomical changes in mental disorders^54^ and associations with cognitive performance^55–57^; with the exception of small associations between brainstem and ADHD^11^, hippocampus and Alzheimer’s disease^12^ and cortical surface area with cognitive function, ADHD, depression and Parkinson’s disease^10^. In general, therefore, there does not appear strong evidence for a significant overlap of common allele influences between cognitive/neuropsychiatric phenotypes and anatomical brain measures. Future research focusing on other brain indices such as white matter microstructure, or using different genetic approaches such as focusing on the genetic overlap at specific loci over genetic correlations across the whole genome^58^, might prove more fruitful.

There are several limitations to our findings. Most noteworthy, the use of a single, homogenously collected and processed UK-Biobank data helps to decrease methodological variation, improving our ability to detect genetic-phenotype associations; however, the UK-Biobank’s cohort does not represent the general UK population, but deviates in important socioeconomic demographics such as age, health, education and economic status^59^, and who we have further limited to only individuals with genetic ancestry of European descent. Moreover, cerebellar measures available from UK-Biobank are created without the use of a cerebellar-specific registration tool, likely leading to poorer registration and segmentation of individuals lobules^60^. For this reason, as well as the high correlation between lobules and its conserved cytoarchitecture, our main analyses focus on total cerebellar volume. We also additionally corrected for potential head motion and position induced artefacts in the scanner to improve the face validity of our results.

In conclusion, we provide a genome-wide association study of the common genetic variation underlying human cerebellar volume. We find, similar to previous reports of cortical and subcortical regions, a moderate-to-high heritability, with generally consistent heritability across the cerebellar lobes. We also report the cerebellum to show the highest genetic similarity to brainstem, pallidal and thalamic volumes, but no significant common allele effect sharing with psychiatric disorders or general cognitive function. While further replication and follow-up functional studies are required, we identify 33 independent SNPs associated with cerebellar volume, highlighting 6 in protein coding variants. Using cerebellar gene expression data, we identify 14 associations that map to eQTLs and 6 associations (4 common with the eQTL-only analysis) showing potential causal relationship with gene expression. In total these additional analyses map associations to 21 unique candidate genes that warrant further investigation. Overall, these results advance our knowledge on the genetic architecture of the cerebellum and pave the way to further research into the neurobiological basis of its anatomy, and associations with normal and abnormal phenotypes.

## Methods

This study used Magnetic Resonance Imaging (MRI) data from the UK-Biobank^15,61^. At the time of initiation of this study in the region of 40,000 individuals’ data had been released. We maintained data separated into two phases containing approximately half of the total sample each, based on our group’s access to the data. We processed and quality controlled each phase independently, compared across phases and then combined the results in a meta-analysis, which we used for all subsequent functional annotation and mapping. Ethics for UK-Biobank was granted by the North West Multi-Centre Ethics Committee, with our study being approved by the UK-Biobank Access Committee

### (Project #17044). Processing genetic data

A full description of UK-Biobank’s data collection, quality control and imputation process can be found elsewhere (http://www.ukbiobank.ac.uk/scientists-3/genetic-data/). Locally, we further harmonised and applied additional quality control (independently) to each phase’s raw genotypes from the UK-Biobank as has been described previously^62^. Briefly, all markers were harmonised to genome build hg19 and common nomenclature based on the Haplotype Reference Consortium r1.1. We excluded markers based on individual marker missingness (>2%), low minor allele count (<5), deviations from Hardy-Weinberg equilibrium (p< 1×10^−10^) and the deviations from the expected Minor Allele Frequency (MAF; >4 standard deviations (SD) from GBR MAF reported in 1000G phase 3). Individuals were removed with excess overall marker missingness rate (>2%) or heterozygosity (>4 × SD from sample mean), those of non-British/Irish ancestry (defined as >4 × SD from 1000G phase 3 GBR sample mean based on first 3 principal components (PCs)) and those with close relatives in the cohort (estimated kinship coefficient > 0.0442 i.e. 3^rd^ degree relatives). Of note, for phase 2 this also included removing individuals with close relatives in phase 1. Of the initial 21,390 and 26,541 individuals with genetic data for phase 1 and phase 2, 19,170 and 22,808 passed our genetic quality control, respectively. From the initial download of over 90M genetic markers, 7,003,604 and 6,935,580 markers remained for phase 1 and phase 2 following quality control, respectively.

### Total cerebellar volume measure generation

We used R(3.6.0) (https://www.R-project.org/) for the generation of our phenotype and all statistical analysis. This study utilises the image derived phenotypes (IDPs) generated from structural T1-weghted MRI scans whose generation and quality control has been described previously^63^. We generated a summated total cerebellar grey-matter volume measure from all the 28 cerebellar lobule IDPs^64^, with the exception of Crus I vermis which was excluded due to its very small size which can cause unreliable results, following previous research^65^. The distribution of cerebellar volume values in each phase were normal. We removed individuals missing any of our key covariates (listed below) and individuals with outlier total cerebellar or total brain grey- and white-matter volume (UK-Biobank data-field code: <u>25010</u>). Outliers were defined as values greater than five times the median absolute deviation from overall median.

To correct for possible imaging-based and other related variables which might confound our result, in a univariate multiple linear regression model we regressed total cerebellar volume on total brain volume, age (UK-Biobank data-field code: <u>21003-2.0</u>), age^2^ (2^nd^ degree orthogonal polynomial), sex (<u>31</u>), age^2^*sex, mean resting-state functional MRI head motion averaged across space and time points (<u>25741-2.0</u>) (log transformed; <u>21001-2.0</u>), imaging centre attended (<u>54-2.0</u>), date attended imaging centre (<u>53-2.0</u>), X-, Y- and Z-head position in the scanner (<u>25756</u>, <u>25757</u>, <u>25758</u>) and starting table-Z position (<u>25759</u>). The residuals derived from this for each phase showed a normal distribution. We scaled the residuals obtained from this model to provide beta’s reflecting changes in standard deviations of residual cerebellar volume.

### Genome-wide association study (GWAS)

Following generation of phenotype measures as outline above, the GWAS for phase 1 included 17,818 participants and for phase 2 15,447 participants (supplementary table 1). Of note the larger drop in phase 2 was explained by the availability of MRI data, rather than differences in quality control filtering between both phases. We removed markers with minor allele counts < 5 within each phase, leaving 6,402,132 and 6,303,745 markers respectively. GWAS analyses were run on PLINK (v1.9)^66^, inputting our cerebellar residuals and covariates of the first 10 genetic PCs to correct for potential effects of remaining population structure. The model assumed linear additive genetic effects. We used LocusZoom^20^ to visually inspect GWAS-significant (p< 5×10^−8^) peaks.

### SNP-based heritability (h^2^_SNP_)

For each phase we estimated the lower-bound of narrow-sense (additive) single nucleotide polymorphism (SNP)-based heritability (h^2^_SNP_) using GCTA-GREML (Genome-wide complex trait analysis – genome-based restricted maximum likelihood)^17,18^ on the raw genotypes. This is done by comparing genetic similarity (in unrelated individuals at our pre-defined cut-off following the above quality control) to phenotypic similarity of our total cerebellar volume measure between participants. As with the GWAS analysis, the first 10 genetic principal components were added to help correct for remaining population structure.

### Identification of independent regions

Genome-wide association signals in each region were refined to identify independently associated signals by applying a stepwise conditional analysis using the COJO (multi-SNP-based conditional & joint association analysis using GWAS summary data) function in GCTA^16,17^. Linkage disequilibrium (LD) data for this analysis was derived from genotypes of the respective UK-Biobank phases. Analysis of correlation structure was limited to 10Mb blocks around genome-wide signals. LD-ranges around index SNPs were defined according to nominally associated LD-partners; specifically, the boundaries around an index SNP defined by modest LD-partner (r^2^> 0.2) with an association of p< 0.05. Additionally, we identified high-LD proxy-SNPs with r^2^> 0.8 to the index SNPs for use in functional annotation.

### Comparison of phase data

#### Replication and two-sided binomial sign test

Independent genome-wide significant association signals in each phase were mapped to GWAS results from the other phase, with replication defined as those passing Bonferroni-corrected significance (p< 0.05/number of index SNPs identified)

#### Genetic Correlation

Genetic correlation (rg) analysis was performed using the LDSC software^67^, regressing the SNP associations (products of the z-scores between the two traits) on their linkage disequilibrium (LD) scores. All summary statistics were limited to a common subset of HapMap3 SNPs prior to analysis. Of note, LDSC regression is not a bounded estimator, therefore, upper bounds of genetic correlation can exceed 1.0 due to sampling variation, though – since none of our results greatly exceeded this level and standard errors were low – we capped them here for display.

#### Polygenic scores

We used PLINK to generate polygenic scores for all participants in each phase, using the summary statistics from the other phase (clumping r^2^> 0.2). We further filtered SNPs at 10 different p-value thresholds: p< 0.5, 0.1, 0.05, 0.01, 0.001, 1×10^−4^, 1×10^−5^, 1×10^−6^, 1×10^−7^ & 1×10^−8^ and repeated this with and without including regions of long-range LD as defined from 1000G phase 3 EUR. Multiple linear regression was used to ascertain the unique variance of total cerebellar volume explained by each polygenic score (ΔR^2^), accounting for the same covariates as used to generate the GWAS (see above section). This was calculated by subtracting the R^2^ of the model without covariates from the R^2^ of model with covariates. Bonferroni correction was applied for the number of tests performed (p< 0.0013 {0.05/(10×2×2)})

### Meta-analysis

We meta-analysed the two phases of GWAS using METAL^19^, weighting the effect size estimates by the inverse of the corresponding standard errors. We retained only the 6,193,476 markers present in both phases. Identification of independent SNPs and calculations of SNP-based heritability were performed using the same methods as outlined above. For the GCTA-GREML analysis of h^2^_SNP_ we created a merged phase dataset using PLINK, so as to obtain the raw genotypes for the whole sample.

### Annotation of GWAS identified independent regions

We annotated associated regions with positional and functional information. Physical annotation of transcripts (ftp://ftp.ensembl.org/pub/grch37/current/gtf/homo_sapiens/Homo_sapiens.GRCh37.87.gtf.gz) was applied using overlap of LD-ranges with transcripts boundaries. Expression quantitative trait loci (eQTL) annotation was based on the GTEx-v7 data (https://gtexportal.org/home) for cerebellum and cerebellar hemisphere labelled tissues, mapped to index and LD-partners. Similarly, index and LD-partner overlap were mapped to SNP consequence (http://www.ensembl.org/), combined annotation-dependent depletion (CADD) Phred-like scores^68^, Polyphen category^69^ and SIFT category^70^.

### Summary-data-based Mendelian randomization (SMR)

We used summary-based Mendelian randomization (SMR)^21,22^ to explore whether the effect size of a SNP on the phenotype is mediated by gene expression. Correlation may infer a causal or pleiotropic relationship – as compared to those caused by linkage - and can prioritise genes within the region for follow-up studies. SMR was implemented using the SMR package (https://cnsgenomics.com/software/smr). The eQTL studies used in the SMR analysis were the same two GTEx-v7 cerebellar labelled tissue data (https://gtexportal.org/home). SMR analysis was limited to genome-wide significant SNPs reported in the cerebellar volume GWAS. To detect heterogeneity of assocations within a region, we applied a HEIDI (heterogeneity in dependent instruments) test, using a conservative threshold (p_HEIDI_≥ 0.05). To provide sufficient data to implement the HEIDI test, analysis was limited to transcripts with a minimum of 10 SNPs in the model. We applied an SMR-wide Bonferroni correction based on the number of transcripts that passed inclusion criteria, for both the cerebellum (p_SMR_< 1.42×10^−6^ {0.05/3526}) and cerebellar hemisphere (p_SMR_< 2.09×10^−5^ {0.05/2389}) labelled tissues.

### Genetic correlation analysis

#### Between study genetic correlation for other brain-related traits

Using the LDSC approach as described above, we calculated genetic correlations between our total cerebellar volume summary statistics and those of other brain-related traits, including previously reported cerebellar measures from Elliott et al (2018)^13^ (FreeSurfer^71^ defined left & right cerebellum and FSL FAST^72^ defined 28 individual cerebellar lobules; n= 8,428 EUR) and Zhao et al (2019)^14^ (ANTs (http://stnava.github.io/ANTs/) defined left & right cerebellar hemispheres and 3 vermal divisions; n= 19,629 EUR). To limit the number of analyses, the comparison with results from Elliott et al were limited to their FreeSurfer analysis. All downloaded summary statistics were harmonised to genome build hg19 and common nomenclature based on the Haplotype Reference Consortium r1.1 and underwent the same procedural steps as outlined above (including HapMap3 filtering_. We also report the LDSC estimated SNP-based heritability scores for the other cerebellar traits, calculated by regressing SNP’s trait association (χ^2^) on their LD. Additionally, to assess the number of novel association regions identified in our meta-GWAS compared to those previously identified in these published works, we deemed novel regions as those with no previously identified index SNP within 500kb of our identified independent regions’ LD ranges or with a previously identified index SNP with r^2^> 0.1 of anyone of our index SNPs.

As several of the identified variants were associated with anthropomorphic measures, in a post-hoc analysis we wished to ascertain that the identified cerebellar variants were generally independent from a collection of anthropomorphic measures collected from the full UK-Biobank cohort (http://www.nealelab.is/uk-biobank/ GWAS round 1 2017 release version limited to EUR ancestry). These included standing height (data-field: <u>50</u>; n= 336,474), sitting height (<u>20015</u>; n= 336,172), birth weight (<u>20022</u>; n= 193,063), body mass index (<u>21001</u>; n= 336,107), weight (<u>21002</u>; n= 336,227) and body fat percentage (<u>23099</u>; n= 331,117).

We also ascertained the genetic correlation with summary statistics of other brain-based measures and brain-related traits. Brain-based measures were those from the ENIGMA group for mean total cortical thickness and surface area using FreeSurfer analysis (n= 33,992 EUR)^10^, and for the hippocampus (n= 26,814 EUR)^12^ and other subcortical volumes of the putamen, pallidum, thalamus, amygdala, nucleus accumbens, caudate nucleus and brainstem (n= 37,741 EUR)^11^. For brain-related psychiatric and neurological traits, we used the latest GWAS summary statistics for schizophrenia (40,675 cases; 64,643 controls)^23^, bipolar disorder (20,352 cases; 31,585 controls)^24^, autism spectrum disorder (18,381 cases; 27,969 controls) (ASD)^25^ and Parkinson’s disease (15,056 cases, 18618 proxies, 430,000 controls)^26^.

Bonferroni correction was used for each set of correlations (cerebellar traits: p< 0.0071 {0.05/7}; anthropomorphic traits: p< 0.0083 {0.05/6}; brain-based traits: p< 0.0050 {0.05/10} & brain-related traits: p< 0.0083 {0.05/6}).

#### Within cerebellum analysis – by lobe analysis

We divided the cerebellum into lobes based on demarcations of primary, horizontal and posterolateral fissures as outlined previously^73^, though grouping hemisphere volumes and separating the flocculonodular lobe. This created 7 lobes, being hemispheres of the anterior (I-V), superior posterior (VI-Crus I), inferior posterior (Crus II-IX) and flocculonodular (X) and separate vermal regions of the latter three (excluding the Crus I vermis). The same outlier exclusion was applied to each lobe separately – as had already been applied to total cerebellar volume – and we removed those individuals with an outlier value (i.e. outside five times median absolute deviation) for any lobe (phase 1: 17,813; phase 2: 15,438; total: 33,251). The same quality control procedures, use of PLINK – along with application of the same covariate list – and METAL analysis were performed for each lobe as done for the main analysis. SNP-based heritability estimates using GCTA-GREML were also similarly obtained. Genetic correlations between lobes, between lobes and other brain-regions, and between lobes and other brain-related traits were calculated using LDSC software using the same procedure as outlined for our primary analysis. Bonferroni adjusted p-values (significance threshold p_Bonferroni_< 0.05) were provided following correction for the number of tests (lobe – lobe correlation: p< 0.00024 {0.05/((7×6)/2}); lobe – other brain regional volume correlation: p< 0.00071 {0.05/(7×10)}; lobe – other trait correlation: p< 0.0012 {0.05/(7×6)})

## Acknowledgements

This research was conducted using the UK-Biobank resource under project ref. 17044 and was supported by the Medical Research Council Programme grant ref. G08005009. TCh was supported by a Wellcome Trust PhD scholarship (ref 203770/Z/16/Z)

## Author Contributions

TCh, XC & RJLA devised the project. TCh, VEP, SL, EB, XC & RJLA participated in imaging and genetic data processing and outputs generation. TCh, RJLA, VEP, SL & EB performed the statistical analyses. TCh, KDS, JTRW, XC & RJLA interpreted the results and wrote the paper. All authors contributed on the discussion of results and revised and approved the final manuscript.

## Competing Interests statement

JTRW has received grant funding from Takeda Pharmaceutical Company for research unrelated to this work. All other authors declare no competing interests.

